# Maternal high-fat/high-sugar diet has short-term dental effects and long-term sex-specific skeletal effects on adult offspring mice

**DOI:** 10.1101/2025.07.06.663396

**Authors:** Mohamed G. Hassan, Kyle Koester, Natalia S. Harasymowicz, Arin K. Oestreich, Kelle H. Moley, Farshid Guilak, Erica L. Scheller

## Abstract

**Background:** Maternal nutrition is increasingly recognized as a modulator of offspring skeletal development. While genetics has long been considered the primary determinant of craniofacial morphology, emerging evidence suggests that prenatal and early postnatal dietary exposures also influence facial morphology. However, how maternal diet differentially affects male and female craniofacial structures remains unclear. This study aimed to examine the effects of a maternal high-fat, high-sugar (HFHS) diet on craniofacial and dental morphology in first-(F1) and second-(F2) generation adult mice.

**Materials and Methods:** Female mice were fed a HFHS diet for six weeks before mating and throughout pregnancy and lactation. F1 offspring were weaned to a standard chow diet, and a subset of female F1 offspring were bred to produce F2 offspring, also maintained on chow. Craniofacial skeletal and dental structures of adult F1 and F2 mice at 1-year of age were assessed using micro-computed tomography for linear and geometric morphometrics.

**Results:** HFHS diet exposure significantly reduced midfacial and mandibular length in F1 females, and these effects persisted in F2 females. Mandibular shape differences were also observed in both generations of females. In males, skull size remained unchanged, though subtle mandibular shape changes were noted in F1 only. Tooth size was reduced in both sexes of F1 offspring but not in F2.

**Conclusion:** Maternal HFHS diet induces sex- and jaw-specific alterations in craniofacial morphology, with skeletal changes persisting in females across generations, while dental effects did not persist beyond one generation. These findings highlight the potential for maternal dietary habits to exert lasting, intergenerational influences on offspring facial form.

## Introduction

The developmental origins of health and disease (DOHaD) concept suggests that environmental exposures during critical developmental windows, such as gestation and lactation, can program lifelong metabolic and musculoskeletal health outcomes in offspring. The western-style diet, characterized by high fat, high sugar, and high caloric density, has emerged as a significant modifier of offspring metabolic and musculoskeletal outcomes.^1^ Metabolic programming, driven by factors such as high-fat (HF) or high-fat/high-sugar (HFHS) diets, induces epigenetic modifications and alters cellular metabolism, predisposing offspring to chronic diseases such as obesity, insulin resistance, and osteoporosis.^1–3^ For example, maternal HFHS diets in mice impair offspring bone quality by reducing trabecular bone volume and cortical thickness in a sex-specific manner, with male offspring showing significant deficits in tibial structure and strength.^2^ Likewise, maternal obesity disrupts skeletal muscle insulin signaling pathways in nonhuman primates, leading to persistent reductions in glucose uptake even in the absence of systemic insulin resistance.^1^ These effects extend intergenerationally, as grandmaternal HFHS exposure in mice alters radial bone geometry in F2 offspring.^2^ Such findings highlight the importance of parental, and in particular, maternal nutrition in shaping offspring metabolic and musculoskeletal health through mechanisms involving mitochondrial dysfunction, epigenetic reprogramming, and tissue-specific response.^4,5^

Craniofacial morphogenesis is a complex process that demands precise coordination of neural crest cell migration, differentiation, and integration with mesodermal tissues to create the bones, cartilage, and soft tissues of the skull and face. Disruptions to the development of the face and skull during pregnancy can give rise to birth defects like cleft lip and palate, craniosynostosis, and other dentoalveolar issues.^6^ These abnormalities can significantly impact a person’s ability to eat, their appearance, and their overall well-being. Maternal diet during pregnancy has been shown to influence offspring craniofacial and dental development, with emerging evidence from animal models linking nutritional intake to skeletal malocclusion and dentofacial anomalies.^7–10^ For example, the modification of calcium-to-phosphorus ratio during pregnancy altered offspring skull morphology in a sex-specific pattern.^8,9^ Additionally, maternal and F1 exposure to high-fat diet (HFD) significantly increased the midface and the palate length of the offspring.^10^ These studies demonstrate the role of maternal dietary intake in modulating offspring craniofacial growth trajectories under varying maternal dietary conditions.

The concept of DOHaD has highlighted how maternal diet can affect the offspring’s craniofacial skeleton.^8,11–15^ However, significant gaps remain in understanding the intergenerational and tissue-specific mechanisms. Current literature focuses mainly on first-generation (F1) offspring. Some studies have shown that maternal diet alteration can lead to persistent changes in F2 skeletal structures, but questions remain about how this phenomena continues to modify craniofacial development and skeletal integrity across generations.^2^ Another factor that can influence developmental programming is the composition of the maternal diets studied. Most research employs high-fat diet models, which may differ from the combined effects of sugar and fat characteristic of Western dietary patterns. This distinction is important, as HFHS diets may uniquely exacerbate developmental disruptions compared to HFD alone. For example, Serirukchutarungsee *et al.* showed that maternal and F1 exposure to HFD changed craniofacial growth in F1 rats, but no studies have looked at how HFHS diets affect craniofacial structures.^10^ The lack of such data limits our understanding of how diet composition influences craniofacial morphogenesis. Finally, sex-specific responses to maternal dietary alterations further complicate our understanding of craniofacial morphogenesis. Evidence suggests a sex-specific response to various alterations in maternal dietary intakes. ^10,9^ These opposite effects highlight the potential interactions between dietary components and sex-specific hormonal or epigenetic regulatory pathways during craniofacial morphogenesis. Such findings show the importance of investigating how maternal diet composition influences sex-specific developmental trajectories and their underlying mechanisms.

This study aimed to examine the long-term effects of maternal and grandmaternal exposure to a high-fat, high-sugar (HFHS) diet on craniofacial structures and dentition in aged offspring. We hypothesized that early maternal exposure to a HFHS diet could have persistent, sex-specific effects on the craniofacial skeleton and dental structures of aged offspring. The findings of this study enhance our understanding of how common lifestyle factors influence craniofacial morphology across the lifespan.

## Material and Methods

### Ethical statement

The Institutional Animal Care and Use Committee at Washington University in St. Louis approved this study. All animal procedures adhered to the National Institutes of Health guidelines for the care and use of laboratory animals.

### Experimental design

Animals were maintained in a temperature-controlled (22 ± 2°C) room under a 12-h light/dark cycle, with ad libitum access to food and water. 10-week-old virgin female C57BL/6J mice were purchased from the Jackson Laboratory. Following a 1-week acclimation period, the mice were randomly assigned to either a control diet group (Pico Laboratory Rodent diet 20; 13% fat, 62% carbohydrates (3.2% sucrose), and 25% protein) or a HFHS group (Test Diet 58R3; 59% fat, 26% carbohydrates (17% sucrose), and 15% protein) for 6 weeks before mating with C57BL/6J control diet fed males. The rationale for the pre-mating dietary intervention was to mimic the sustained exposure to Westernized diet common in human populations, thus increasing the translational relevance of the study. Pregnant dams continued their assigned diets throughout pregnancy and lactation as previously described.^2^ At 3 weeks of age, the male and female offspring (F1) were weaned and transitioned to a standard chow diet, housed with same-sex animals from their respective maternal treatment groups, and raised until they reached 1 year of age. At the study endpoint, the mice were humanely euthanized via CO_2_ asphyxiation followed by cervical dislocation. To generate F2 offspring, female F1 mice from each maternal dietary treatment group were mated with chow-fed C57BL/6J male mice at 8–10 weeks of age. The resulting F2 offspring were maintained on a standard chow diet and evaluated at 1 year of age (**Figure 1**).

**Figure 1.**
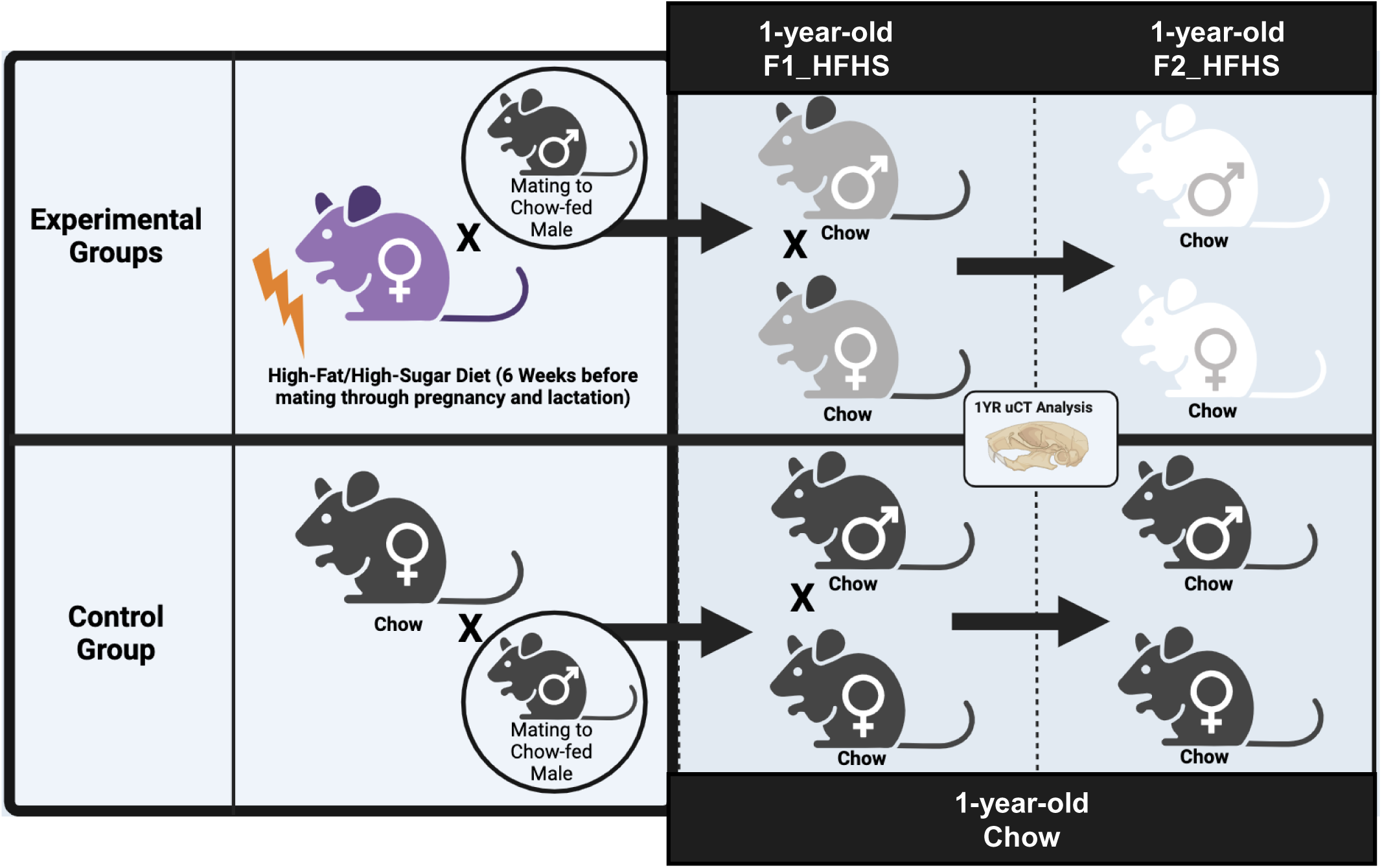
Graphical overview of the study design. The schematic illustrates the overall workflow and experimental design of the study. Timelines, experimental groups (e.g., Chow, F1-HFHS, F2-HFHS), and analytical techniques are represented to summarize the stepwise approach used to address the study objectives.

### Micro-computed Tomography and image acquisition

Dissected heads were fixed in 10% formalin for 48 hours at room temperature. After fixation, specimens were thoroughly rinsed with 70% ethanol and stored at 4°C until scanning. The heads were scanned using a Scanco µCT 50 in the Musculoskeletal Research Center Core at Washington University School of Medicine in St. Louis, United States. Specimens were carefully positioned within the scanner tube to optimize image acquisition and scanned at 17-micron resolution (55kV, 145mA, 0.5mm Al filter). The raw micro-CT data was reconstructed into a three-dimensional volume using Scanco Medical’s μCT Evaluation software. Following reconstruction, the data was exported as DICOM files and imported into Dragonfly software (v4.1) for visualization and analysis.

### Dental measurements

For the dental analyses, radiographic measurements were obtained for the right upper and lower first molars and incisors. To measure the molars, CT cross-sections were centered in the middle of the upper or lower right first molar, with the cross-sections aligned along the anteroposterior axis of the molars, the central pulp, and the molar root. For the incisors, cross-sections were centered in the middle of the lower right first molar but aligned with the anteroposterior axis of the incisor (**Supplemental Figure 1**). The following measurements were taken using Dragonfly software (v4.1): anteroposterior molar length, molar surface area, root length, and incisor length. The selection of these parameters was based on their relevance in characterizing the overall size and shape of the dental structures.^8^

### Skeletal linear measurements

Craniofacial skeletal variation was initially analyzed using traditional linear morphometrics. The following measurements were recorded: skull length, midfacial length, skull width, neurocranium width, maxillary alveolar bone length (right and left), maxillary intermolar width (anterior and posterior), mandible length (right and left, both condylar and gonial), mandible height (right and left), mandibular alveolar bone length (right and left), mandibular intermolar width (anterior and posterior), inter-condylar width, and inter-gonial width (**Supplemental Table 1**). ^16^ All bilateral measurements were averaged between the right and left sides and performed using 3D slicer software.

### Geometric morphometric analysis

Three-dimensional coordinates of 44 cranial and 13 paired mandibular landmarks were collected using Landmark Editor software (**Supplemental Table 2-3**). All landmarking was performed by a single observer (K.K.), who was blinded to group assignments during data collection to minimize bias. Landmark coordinates were exported as text files and imported into *MorphoJ* software for geometric morphometric analysis.^17^ To remove variation due to position, orientation, and scale, a generalized Procrustes analysis (GPA) was performed separately for cranial and mandibular datasets. The aligned Procrustes coordinates were then subjected to principal component analysis (PCA) to summarize the major axes of shape variation. Principal component (PC) plots and wireframe deformation graphs were generated to visualize differences in craniofacial shape among groups.

### Statistical analysis

Descriptive statistics were calculated for skeletal and dental linear measurements. Group differences were assessed using one-way ANOVA, with statistical significance set at *p* < 0.05. GraphPad Prism (Version 10.0.0) was used to perform tests and generate graphics. Intra-observer reliability for skeletal and dental measurements was calculated by repeating measurements on 10 randomly selected samples, with a one-week interval between sessions. Intraclass Correlation Coefficient (ICC), Paired *t*-tests and Bland–Altman plots were used to assess consistency, and all measurements were found to be highly reliable (**Supplemental Table 4).** For landmark reproducibility, the same observer (K.K.) placed 3D landmarks on 7 randomly selected specimens twice, spaced one week apart. Landmark reliability was confirmed by visual inspection of Procrustes distances and by comparing repeated configurations. (**Supplemental Figure 2**)

## Results

Data were analyzed in three groups: chow controls, first generation offspring from HFHS fed mothers (F1_HFHS), and second-generation offspring from HFHS fed grandmothers (F2_HFHS) all at 1-year of age. The total sample size (n= 73) for each group was as follows: F1_Chow (females, *n* = 19; males, *n* = 15), F1_HFHS (females, *n* = 9; males, *n* = 5), and F2_HFHS (females, *n* = 11; males, *n* = 14). The results were organized by both the type of dietary intervention and the sex of the mice. Linear and geometric morphometric approaches were employed to evaluate morphological variation, each addressing different aspects of form. Linear measurements quantified size changes, offering precise and reproducible data on key dimensions including length, width, and height—essential for assessing growth and volumetric differences.^18^ However, as linear morphometrics alone cannot capture complex morphological variation, geometric morphometric analysis was also conducted. This approach utilizes the spatial configuration of anatomical landmarks to assess shape variation independently of size, orientation, and position.^19^ Together, these methods enabled a detailed characterization of craniofacial morphology, distinguishing true shape changes from size-related variation. Specifically, 3D linear measurements were used to assess changes in dental, cranial, and mandibular size, while geometric morphometric analysis—based on landmark placement on the cranium and mandible—evaluated shape changes in these regions.

### Sex-specific skeletal changes persist across F1 and F2 generations

Cranial dimensions showed specific changes in midface length only among female offspring across both generations. Specifically, F1_HFHS and F2_HFHS females exhibited significantly shorter midface length compared to controls, whereas male offspring in both generations did not display any significant changes in cranial dimensions (**Figure 2A-D**, **2A’-D’, Supplemental Figure 3)**. Principal component analysis (PCA) of cranial landmarks didn’t identify a distinct pattern associated with either sex or generation. The first principal component (PC1) accounted for around 13% of the variance and effectively separated males from females, with males clustering towards negative PC scores. PC3, explaining 9.2% of the variance, distinguished Chow and F2_HFHS generations. However, no significant differences were found in overall cranial shape between F1_HFHS, F2_HFHS, and control groups for either sex. Morphological shape changes were not identifiable in 3D cranial wireframes (**Figure 2E-G**).

**Figure 2.**
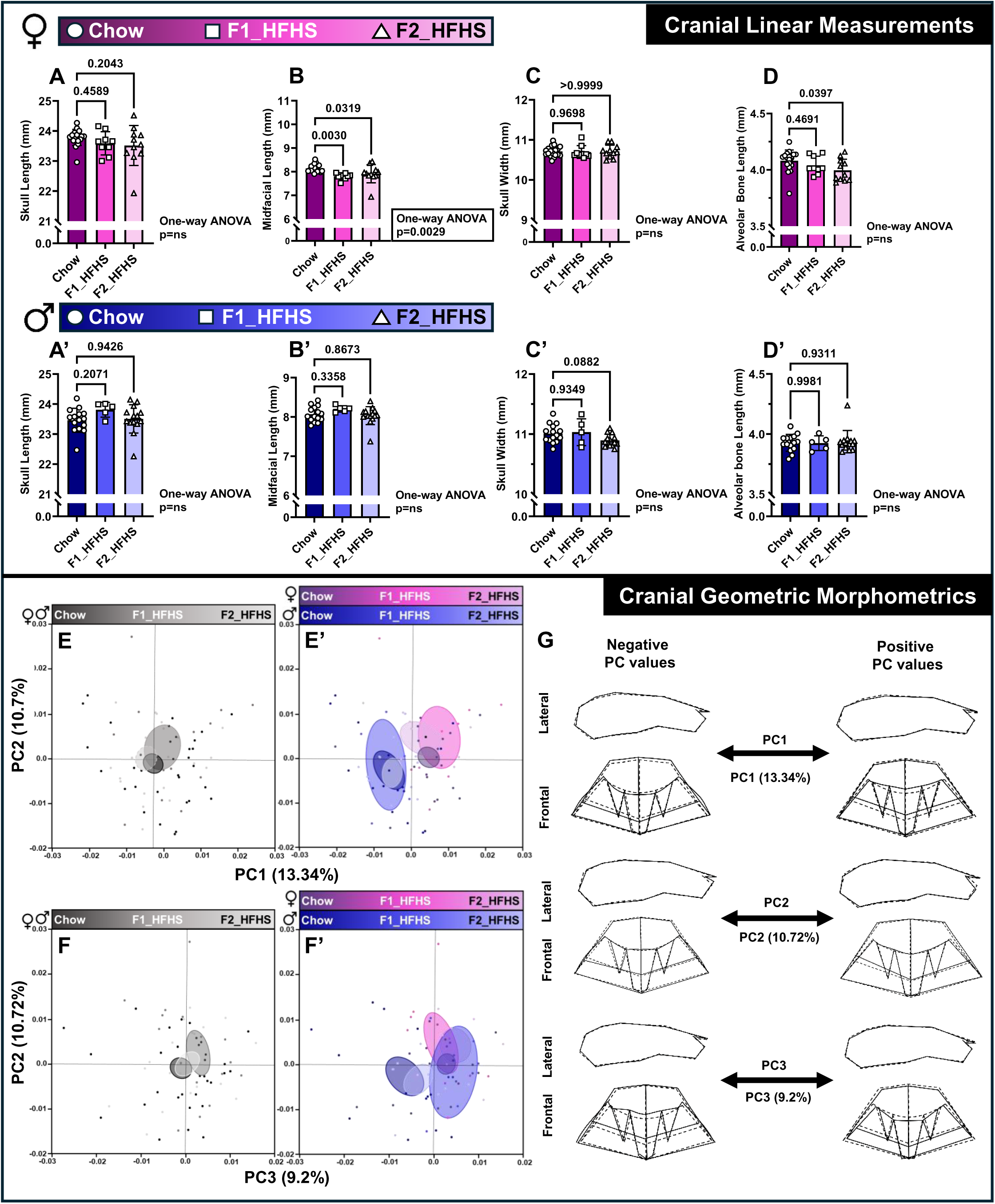
Cranial size and shape changes across experimental groups. A–D shows cranial linear measurements in female offspring, while panels A′–D′ show the corresponding measurements in males. Groups include Chow, F1-HFHS, and F2-HFHS. Data were analyzed using one-way ANOVA followed by pairwise comparisons for significant results. Black rectangles highlight measurements with significant ANOVA outcomes. Female groups are shaded in a pink gradient, and male groups in a blue gradient. E and F represent the first three principal components (PCs) of cranial geometric morphometric analysis, with clustering based solely on intervention. Corresponding panels E′ and F′ show clustering based on both gender and intervention. G illustrates wireframes of shape variation along the extreme ends of PC scores; solid lines represent the shape at the extreme positive and negative PC values, while dotted lines indicate the mean cranial shape.

For mandible, linear morphometrics revealed that F1_HFHS and F2_HFHS females exhibited a significant reduction in the anteroposterior dimension (**Figure 3A-D, Supplemental Figure 4**), as measured by decreased condylar and gonial mandibular lengths (**Figure 3A-D, Supplemental figure 4**). These reductions were positively correlated with midfacial shortening observed in the cranium of the same female cohorts (**Supplemental Figure 6**). In contrast, no significant mandibular differences were detected in male offspring across generations, except only in the inter-molar width (**Figure 3A’-D’, Supplemental Figure 4)**. Geometric morphometric analysis further revealed more robust changes in mandibular shape compared to the cranium. When size effects were removed, principal component analysis showed that PC1 to PC3 explained the majority of the shape variation (55% of total). PC1, accounting for ∼38% of total shape variance, primarily reflected changes in the anteroposterior dimension of the mandible and distinguished F1_HFHS offspring from Chow (**Figure 3E**). PC2, which explained an additional ∼16% of variance, was associated with sexual dimorphism and separated male and female offspring across groups (**Figure 3E’**, **3F’**). These findings suggest that maternal HFHS exposure induces alterations in mandibular shape that are distinct from sex-based morphological variation. F1_HFHS offspring displayed shape changes characterized by shortened mandibular length, reduced height and width of the mandibular ramus, and transverse widening (**Figure 3G**).

**Figure 3.**
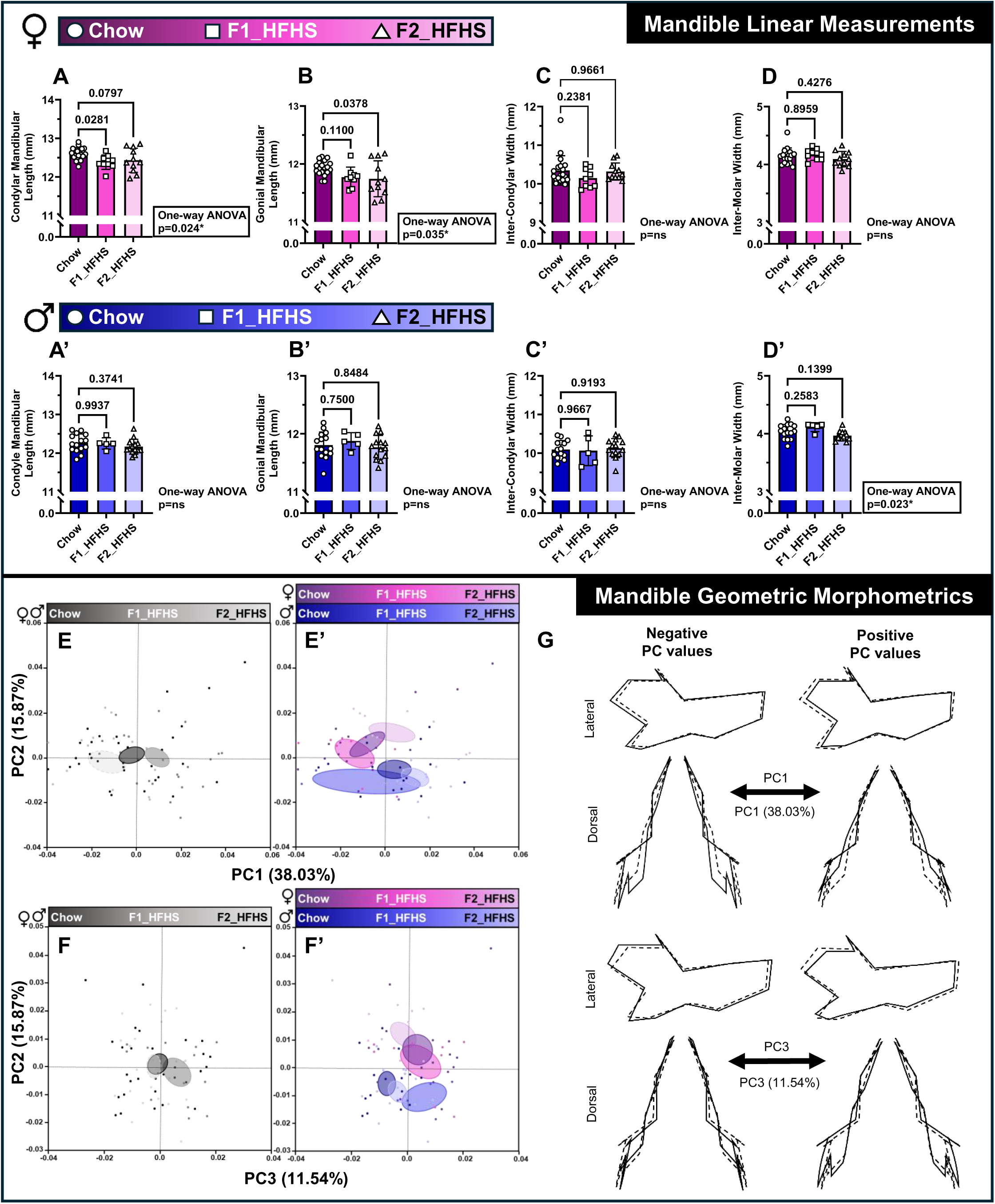
Mandible shape and size changes across experimental groups. A–D shows mandibular linear measurements in female offspring, while panels A′–D′ show the corresponding measurements in males. Groups include Chow, F1-HFHS, and F2-HFHS. Data were analyzed using one-way ANOVA followed by pairwise comparisons for significant results. Black rectangles highlight measurements with significant ANOVA outcomes. Female groups are shaded in a pink gradient, and male groups in a blue gradient. E and F represent the first three principal components (PCs) of cranial geometric morphometric analysis, with clustering based solely on intervention. Corresponding panels E′ and F′ show clustering based on both gender and intervention. G illustrates wireframes of shape variation along the extreme ends of PC scores; solid lines represent the shape at the extreme positive and negative PC values, while dotted lines indicate the mean cranial shape.

### Short-term dental changes occur only in F1 generation in both sexes

In the F1 generation, both males and females born to HFHS diet-fed mothers showed a significant reduction in several upper and lower dental measurements compared to the control group (**Figure 4**). Specifically, the molar anteroposterior length and crown surface area were significantly reduced in the F1_HFHS male and female offspring, affecting both the upper and lower dentition, as well as in the upper molar of the F1_HFHS male offspring (**Figure 4A’**, **4C, Supplemental Figure 5**). Additionally, the root length of female F1_HFHS offspring lower molar showed a significant decrease (**Figure 4D**). Although a decrease was observed in the mandibular incisor length of F1_HFHS offspring, this change was not statistically significant (**Supplemental Figure 5**). The decrease of molar dimensions was not correlated with the maxillary and mandibular alveolar bone length (**Supplemental Figure 6**). Finally, no significant differences were seen in the dental measurements between the F2_HFHS mice and the controls regardless of sex, indicating that the reduction seen in the F1_HFHS generation did not persist in the subsequent generations.

**Figure 4.**
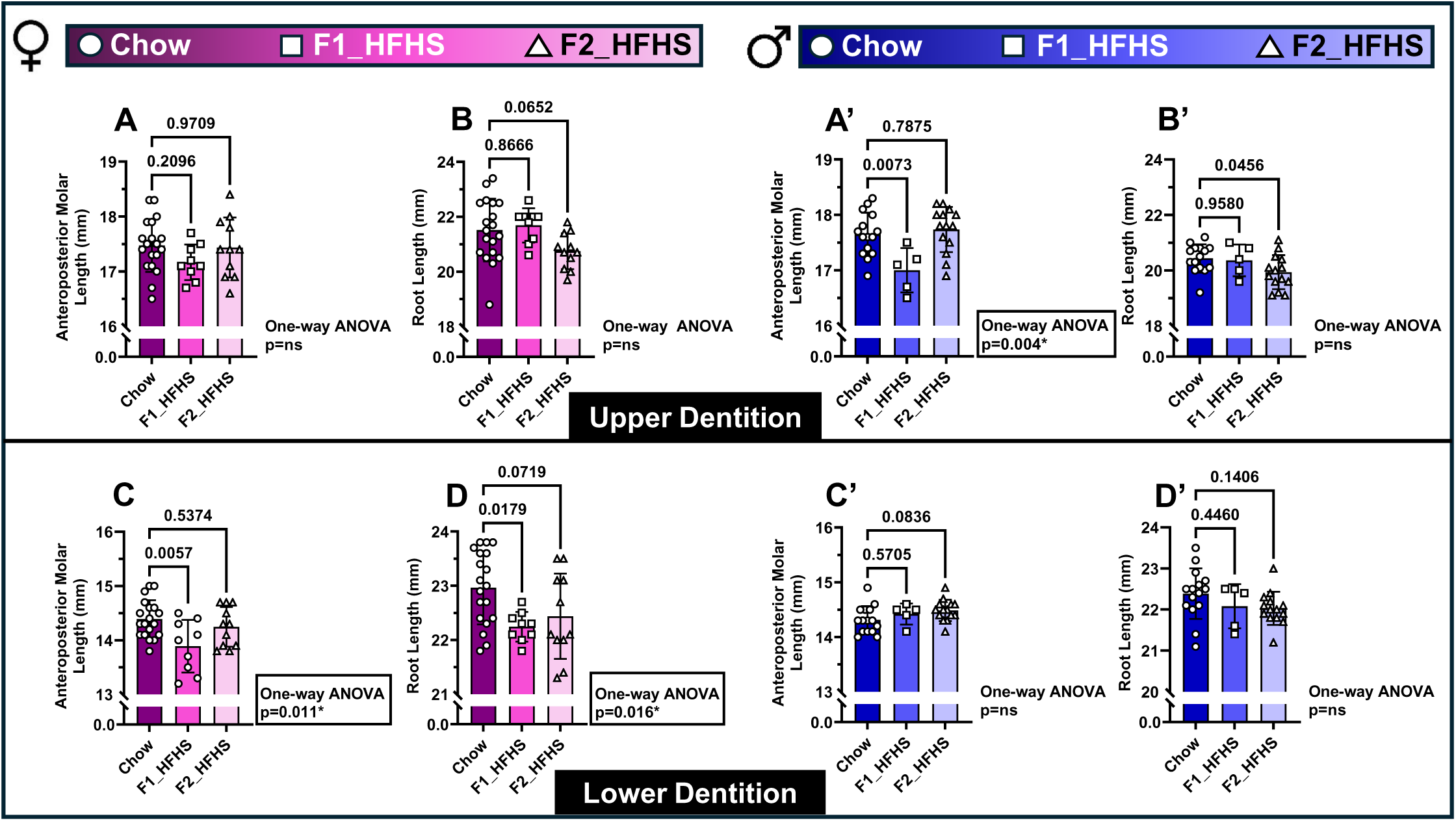
Dental changes across experimental groups. A–D display upper and lower dental measurements in female offspring, while panels A′–D′ show the corresponding measurements in males. Groups include Chow, F1-HFHS, and F2-HFHS. Data were analyzed using one-way ANOVA followed by pairwise comparisons for significant results. Black rectangles highlight measurements with significant ANOVA outcomes. Female groups are shaded in a pink gradient, and male groups in a blue gradient.

## Discussion

The findings of this study show that a maternal HFHS diet induces sex-specific effects on the craniofacial skeleton and dentition of offspring, with skeletal changes in females persisting across generations. Together with our previous studies,^2^ our results provide comprehensive evidence that early nutritional exposures can differentially shape skeletal development in male and female offspring, supporting the concept of DOHaD. Interestingly, the female offspring displayed persistent midfacial and mandibular shortening across generations, while the dental alterations were confined to the first generation, affecting both females and males. These results add a novel dimension to our understanding of how maternal diet composition, particularly HFHS diet, can influence craniofacial growth trajectories across the lifespan and potentially across generations. A previous study in young Japanese women demonstrated that nutritional status, particularly a high-calorie diet pattern, was significantly associated with facial morphology, specifically midfacial retrusion identified through multivariate analyses.^20^ These findings align with our observations, in which female offspring exposed to a maternal HFHS diet exhibited persistent midfacial deficiency into adulthood.

Maternal nutrition is a critical factor in fetal development, with far-reaching implications for craniofacial structures of the offspring. The complex interplay between maternal dietary components and fetal craniofacial development represents an essential area of research, with significant clinical relevance for the early prevention of malocclusions that can arise in settings of compromised skeletal development.^21,22^ Previous studies demonstrate that maternal diet impacts offspring craniofacial development through various nutritional pathways and molecular mechanisms.^9,13–15,20^ These dietary influences can lead to lasting structural alterations that persist long into postnatal life. Unlike previous studies that evaluated younger offspring at 6- to 10-weeks-old,^9,10^ we analyzed offspring at 1-year of age, similar to age 30 years in humans.^23^ This helps us to determine whether early craniofacial changes reflected persistent skeletal adaptations or merely transient developmental alterations. Our findings reveal that skeletal changes, specifically midfacial and mandibular shortening, lasted into adulthood, but only in female offspring. In contrast, the dental changes were limited to the first generation and were observed in both males and females, suggesting a more transient and potentially reversible phenotype.

The development and expression of sexual dimorphism in the craniofacial skeleton are influenced by multiple factors, including genetic programming, hormonal influences, environmental factors, and nutrition. While direct evidence of maternal dietary effects on sex-specific craniofacial development in offspring is limited, previous studies have shown that maternal diet can induce sex-specific changes in the offspring craniofacial skeleton. ^8–10^ Several biological mechanisms related to developmental programming and tissue-specific responses can account for the skeletal changes in female offspring that persist into adulthood, while dental changes affect both sexes but remain limited to the first generation. Previous studies showed that maternal HFD may reduce adult offspring bone mineral density (BMD) and bone volume while delaying skeletal development due to modifications in the epigenetic regulation of osteoblasts that promote cellular senescence to suppress bone formation.^24^ In addition to the potential effect on osteoblasts, a HFHS diet has been shown to increase bone fragility by altering osteocyte function.^25^ Sex hormones also significantly influence bone remodeling throughout life.^26^ Maternal HFHS diets have been shown to alter sex-hormone levels in offspring, which could disproportionately affect female skeletal development through estrogen-dependent pathways.^27^ This may explain why skeletal changes persist specifically in female offspring.

The developmental timing of dietary exposure is a likely mechanism explaining why craniofacial phenotypes observed in the F1 generation persist in the F2 generation. Although the HFHS diet is not directly given to F1 offspring, the oocytes of F1 offspring mice are exposed to HFHS during the pregnancy of the mother. These F1 germ cells may thus undergo significant epigenetic reprogramming events that could impact the F2 offspring, despite the F1 parent not being fed HFHS diet during pregnancy or lactation. This mechanism is supported by evidence indicating that maternal diet during gestation can induce epigenetic changed of the oocytes, leading to long-term effects on subsequent offspring.^28^ Furthermore, studies have shown that maternal HFD can induce epigenetic modifications in oocytes; however, these modifications may not persist into F2 and F3 generations without continued environmental pressure.^29^ These findings collectively suggest that direct exposure during F1 oocyte development is a key mechanism driving persistent F2 phenotypes, while germline reprogramming may buffer or erase these effects in subsequent generations.

Improved understanding of how environmental factors, such as diet, influence craniofacial morphology across generations may offer valuable insights for preventing and intervening early in the development of malocclusion. Previous studies demonstrated that prolonged consumption of a soft diet across multiple generations can lead to long-term skeletal changes in adult mice.^16^ While existing studies have focused on the craniofacial morphological alterations associated with dietary shifts, the underlying mechanisms driving these changes remain unclear. Clinically, changes in patient demographics and lifestyle—particularly dietary habits—warrant closer consideration to support more personalized treatment strategies.

Lifestyle-driven changes in growth trajectories may require orthodontic interventions to be tailored not only to developmental timing but also to sex-specific skeletal responses. Future studies are needed to understand these mechanistic pathways to develop biologically based, individualized treatment plans for malocclusion prevention and treatment.

A limitation of the present study is the unequal sample distribution across groups, particularly when stratified by treatment and sex. This challenge arises from the inherent nature of the longitudinal in-vivo studies. Maintaining rigorous control over breeding outcomes throughout an extended period of over three years is difficult, as fluctuations in litter size, sex ratios, and offspring survival can impact group size. While some variation is normal in biological studies, future work could use larger groups or breeding methods to overcome these problems.

## Conclusion

In conclusion, it is important to consider that environmental factors, such as maternal diet, mechanical forces, and metabolic conditions, can interact with key developmental pathways, ultimately shaping craniofacial morphology. The findings of this study show that maternal intake of a HFHS diet induces sex- and jaw-specific alterations in craniofacial morphology, with female offspring displaying persistent reductions in midfacial and mandibular length across F1 and F2 generations. By contrast, dental changes were more transient, affecting both males and females in the F1 generation only. Further research is needed to understand the mechanisms behind these observed changes, which may provide insights into the environmental contributors to malocclusion.

