## Supplemental File for "Maternal high-fat/high-sugar diet has short-term dental effects and long-term sex-specific skeletal effects on adult offspring mice"

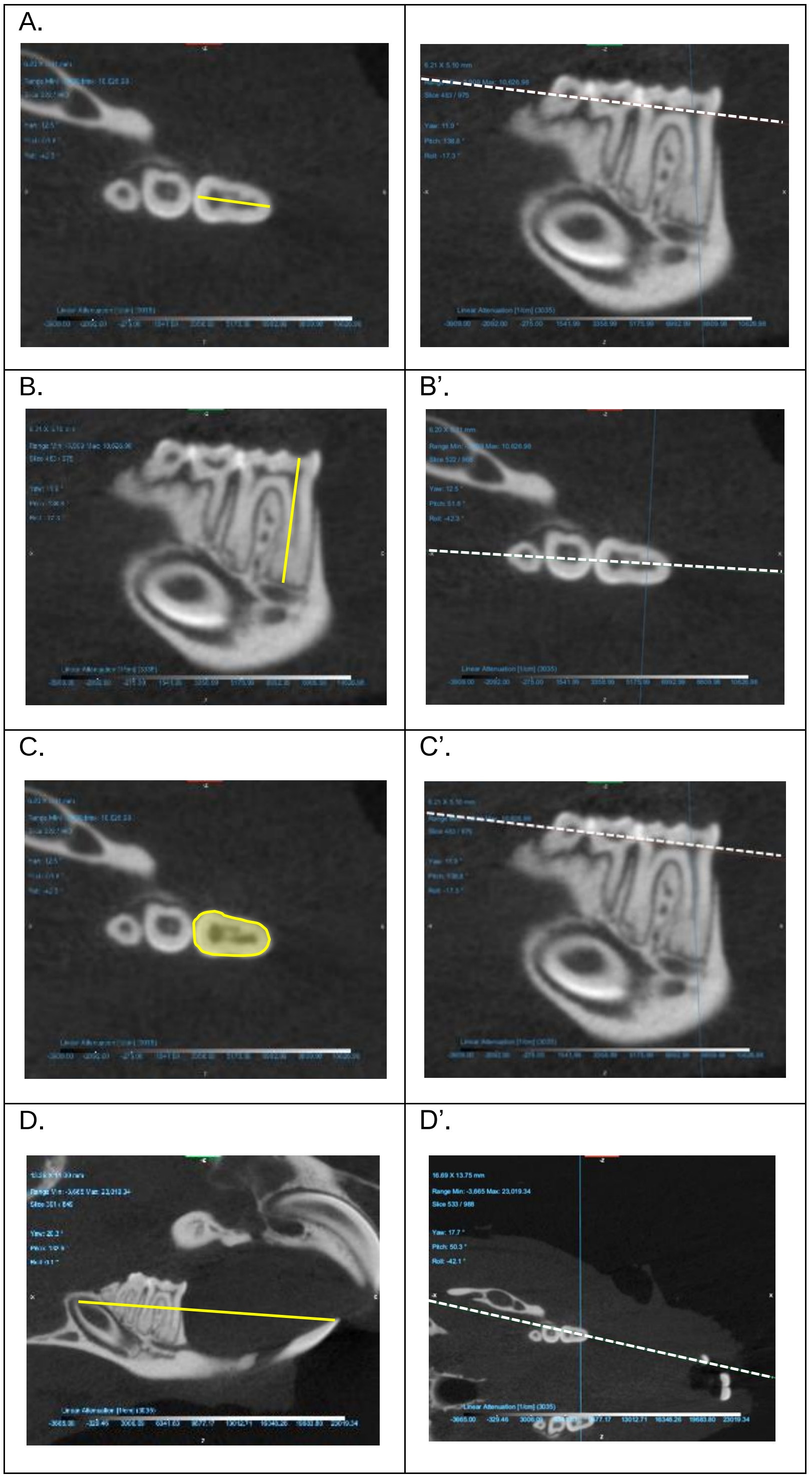


**Supplemental Figure 1. Dental Measurements and Axis Alignment.** (A) Anteroposterior length of the first molar, (B) Root length of the first molar, (C) Cross-sectional surface area of the first molar, (D) Anteroposterior length of the incisor. (A'–D') Corresponding axial sections used to obtain each measurement shown in (A–D), respectively. Yellow solid lines indicate the collected measurement, while white dotted lines indicate the anatomical viewing plane.

**Supplemental Table 1**

**3D Skeletal Linear Measurements.** This table summarizes the linear skeletal measurements extracted from 3D reconstructions. The left column lists the numerical identifiers assigned to each measurement. The middle column provides a detailed anatomical description corresponding to each measurement. The right column includes visual representations illustrating the location and orientation of each linear measurement on the cranium and mandible.

| | | |
|---|---|---|
| 1 | Skull Length | 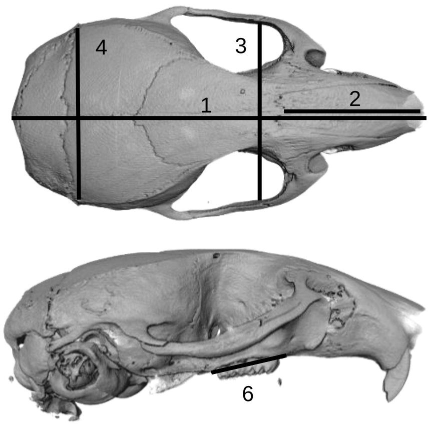 |
| 2 | Midfacial Length | |
| 3 | Skull Width | |
| 4 | Neurocranium Width | |
| 5 | Upper Lt. Molar Alveolar Bone | |
| 6 | Upper Rt. Molar Alveolar Bone | |
| 7 | Upper Anterior Intermolar Width | |
| 8 | Upper Posterior Intermolar Width | |
| 9 | Rt. Mandibular Length Condyle | 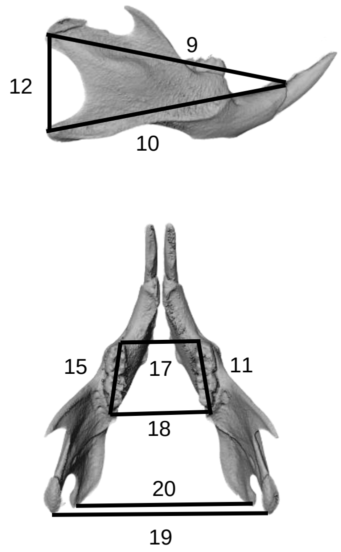 |
| 10 | Rt. Mandibular Length Gonial | |
| 11 | Lower Rt. Molar Alveolar Bone | |
| 12 | Rt. Mandibular Height | |
| 13 | Lt. Mandibular Length Condyle | |
| 14 | Lt. Mandibular Length Gonial | |
| 15 | Lower Lt. Molar Alveolar Bone | |
| 16 | Lt. Mandibular Height | |
| 17 | Lower Anterior Intermolar Width | |
| 18 | Lower Posterior Intermolar Width | |
| 19 | Intercondylar Width | |
| 20 | Intergonial Width | |

**Supplemental Table 2**

**Cranium Landmarks and anatomical Locations.** This table presents the cranial landmarks used for morphometric analysis. The left column lists the assigned landmark numbers, while the right column provides detailed anatomical descriptions corresponding to each landmark. Visual references depicting the location of each landmark on the cranium are displayed in the bottom row of the table to aid in spatial orientation and reproducibility.

| | |
|---|---|
| 1 | Most anterior suture of nasal bone |
| 2 | Most posterior suture of nasal bone |
| 3 | Most posterior suture of frontal bone |
| 4 | Most posterior suture of parietal bone |
| 5 | Most posterior point on the median line of interparietal bone |
| 6 & 7 | Right and Left most anterior point of suture between frontal and parietal bones |
| 8 & 9 | R. and L. intersection between parietal, occipital and squamosal bones |
| 10 & 11 | R. and L. most posterior junction of squamosal bone and zygomatic process |
| 12 & 13 | R. and L. most anterior suture of jugal bone and the zygomatic process of the maxillary bone |
| 14 & 15 | R. and L. zygomaticomaxillary suture |
| 16 & 17 | R. and L. intersection of the frontal, lacrimal, and zygomatic process of the maxillary bone |
| 18 & 19 | R. and L. infraorbital foramen most superior part |
| 20 & 21 | R. and L. premaxilla-right nasal bone most anterior point of suture / bottom of nasal foramen |
| 22 & 23 | R. and L. most superior part of incisor alveolus |
| 24 & 25 | R. and L. most inferior anterior point of incisor alveolus |
| 26 & 27 | R. and L. most inferior posterior point of incisor alveolus |
| 28 & 29 | R. and L. premaxilla- maxilla most ventral junction |
| 30 & 31 | R. and L. most anterior point of first molar alveolus |
| 32 & 33 | R. and L. most posterior point of third molar alveolus |
| 34 & 35 | R. and L. most anterior point of palatine foramen |
| 36 & 37 | R. and L. most posterior point of palatine foramen |
| 38 & 39 | R. and L. most posterior point of hamular process |
| 40 | Midline point of suture between occipital and basisphenoid bones |
| 41 | Midline point of basisphenoid and presphenoid bones |
| 42 | Midline point of suture between occipital and sphenoid bones |
| 43 | Most anterior point of foramen magnum, basion |
| 44 | Most posterior point of foramen magnum, bregma |

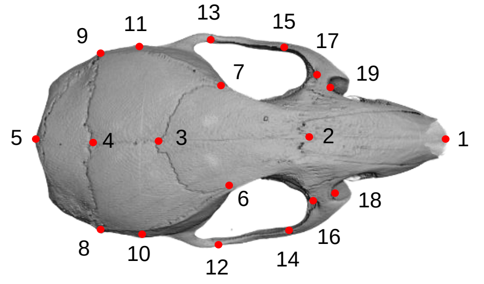


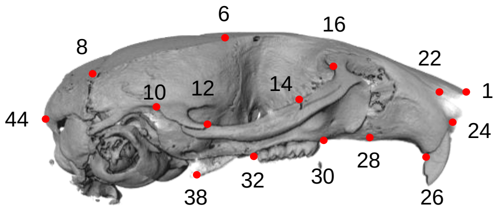

**Supplemental Table 3**

**Mandible Landmarks and anatomical Locations.** This table outlines the anatomical landmarks used for mandibular morphometric analysis. Landmark numbers are listed in the left column, with corresponding anatomical descriptions provided in the middle column. The right column includes visual depictions of each landmark plotted on the mandible, facilitating accurate identification and spatial orientation.

| | | |
|---|---|---|
| 1 & 2 | Most superior point of the incisor alveolus | 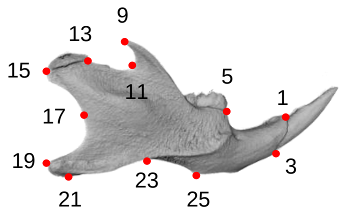  |
| 3 & 4 | Most inferior point of the incisor alveolus | |
| 5 & 6 | Most anterior point of the first molar alveolus | |
| 7 & 8 | Most posterior point of the third molar alveolus | |
| 9 & 10 | Most posterior tip of the coronoid process | |
| 11 & 12 | Most concave point of the coronoid process | |
| 13 & 14 | Most anterior point of the articular surface of the condyle | 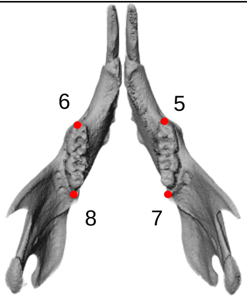  |
| 15 & 16 | Most posterior tip of the condyle | |
| 17 & 18 | Most concave point between condyle and gonial | |
| 19 & 20 | Most posterior tip of the gonial angle | |
| 21 & 22 | Most inferior point of the gonial angle | |
| 23 & 24 | Ascending ramus dorsal-most ventral point | |
| 25 & 26 | Alveolar region most inferior point | |

**Supplemental Figure 2**

**Example of the intra-reliability test demonstrating the consistency of landmarking in the cranium and mandible**

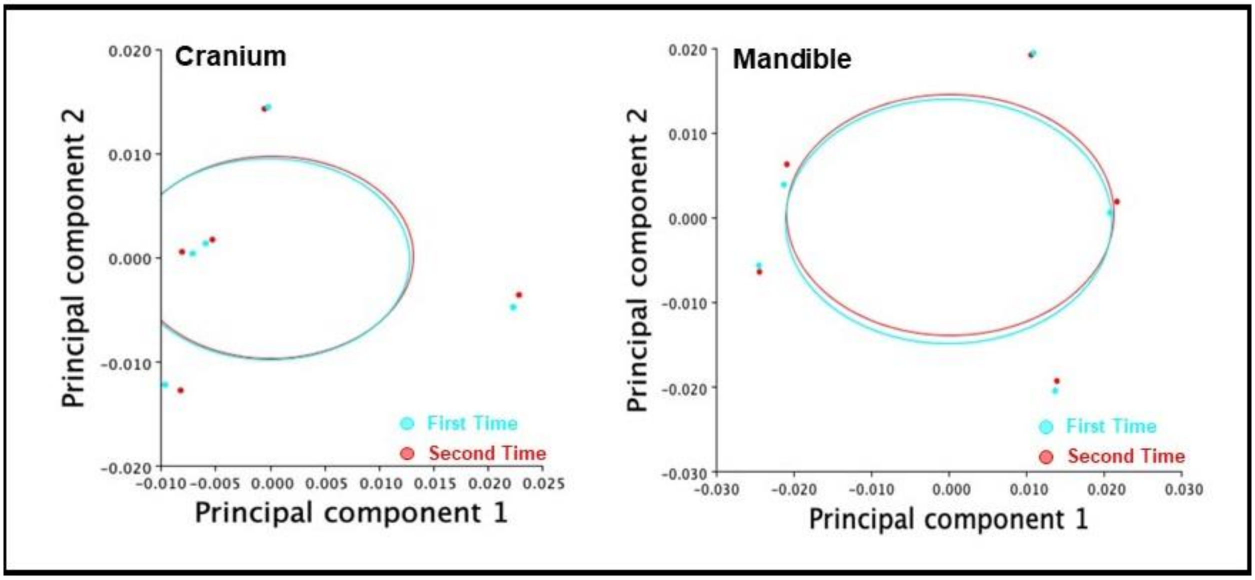

### Supplemental Table 4

### ICC used to determine the intra-observer reliability of the repeated measurement errors for linear measurements

| | Intraclass Correlation | 95% Confidence Interval | | | |
|---|---|---|---|---|---|
| | | Lower Bound | Upper Bound | Value | Sig |
| **Skull Length** | | | | | |
| Single Measures | .99 | 0.98 | 1 | 389 | 0.000 |
| Average Measures | 1.00 | 0.99 | 1 | 389 | 0.000 |
| **Midfacial Length** | | | | | |
| Single Measures | .99 | 0.95 | 1 | 157 | 0.000 |
| Average Measures | .99 | 0.98 | 1 | 157 | 0.000 |
| **Skull Width** | | | | | |
| Single Measures | .89 | 0.63 | 0.97 | 17 | 0.000 |
| Average Measures | .94 | 0.78 | 0.98 | 17 | 0.000 |
| **Neurocranium Width** | | | | | |
| Single Measures | .96 | 0.84 | 0.99 | 44 | 0.000 |
| Average Measures | .98 | 0.91 | 0.99 | 44 | 0.000 |
| **Left Alveolar Length (Mandible)** | | | | | |
| Single Measures | .92 | 0.71 | 0.98 | 23 | 0.000 |
| Average Measures | .96 | 0.83 | 0.99 | 23 | 0.000 |
| **Right Alveolar Length (Mandible)** | | | | | |
| Single Measures | .82 | 0.45 | 0.93 | 10.6 | 0.001 |
| Average Measures | .90 | 0.62 | 0.97 | 10.6 | 0.001 |
| **Average Alveolar Length (Mandible)** | | | | | |
| Single Measures | .96 | 0.86 | 0.99 | 9 | 0.000 |
| Average Measures | .98 | 0.93 | 1.00 | 9 | 0.000 |
| **Anterior IMW (Mandible)** | | | | | |
| Single Measures | .84 | 0.50 | 0.96 | 11 | 0.001 |
| Average Measures | .91 | 0.66 | 0.98 | 11 | 0.001 |
| **Posterior IMW (Mandible)** | | | | | |
| Single Measures | .94 | 0.79 | 0.98 | 32 | 0.000 |
| Average Measures | .97 | 0.88 | 0.99 | 32 | 0.000 |
| **Right Mandible Length (Condyle)** | | | | | |
| Single Measures | .91 | 0.49 | 0.98 | 35 | 0.000 |
| Average Measures | .95 | 0.66 | 0.99 | 35 | 0.000 |
| **Right Mandible Length (Gonial)** | | | | | |
| Single Measures | .84 | 0.28 | 0.96 | 20 | 0.000 |
| Average Measures | .92 | 0.44 | 0.98 | 20 | 0.000 |
| **Right Alveolar Length (Cranium)** | | | | | |
| Single Measures | .95 | 0.82 | 0.99 | 41 | 0.000 |
| Average Measures | .97 | 0.90 | 0.99 | 41 | 0.000 |
| **Right Mandibular Height** | | | | | |
| Single Measures | .98 | 0.94 | 1 | 130 | 0.000 |
| Average Measures | .99 | 0.97 | 1 | 130 | 0.000 |
| **Left Mandibular Length (Condyle)** | | | | | |
| Single Measures | .98 | 0.94 | 1 | 113 | 0.000 |
| Average Measures | .99 | 0.97 | 1 | 113 | 0.000 |
| **Left Mandibular Length (Gonial)** | | | | | |
| Single Measures | .96 | 0.87 | 0.99 | 55 | 0.000 |
| Average Measures | .98 | 0.93 | 1.00 | 55 | 0.000 |
| **Left Alveolar Length (Cranium)** | | | | | |
| Single Measures | .96 | 0.84 | 0.99 | 45 | 0.000 |
| Average Measures | .98 | 0.92 | 0.99 | 45 | 0.000 |
| **Left Mandibular Height** | | | | | |
| Single Measures | .95 | 0.83 | 0.99 | 40 | 0.000 |
| Average Measures | .98 | 0.91 | 0.99 | 40 | 0.000 |
| **Anterior IMW (Cranium)** | | | | | |
| Single Measures | .76 | 0.32 | 0.93 | 7.4 | 0.003 |

| | | | | | |
|---|---|---|---|---|---|
| Average Measures | .86 | 0.59 | 0.97 | 7.4 | 0.003 |
| **Posterior IMW (Cranium)** | | | | | |
| Single Measures | .88 | 0.55 | 0.97 | 20 | 0.000 |
| Average Measures | .94 | 0.71 | 0.98 | 20 | 0.000 |
| **Inter-Condylar Width** | | | | | |
| Single Measures | .89 | 0.63 | 0.97 | 17 | 0.000 |
| Average Measures | .94 | 0.77 | 0.98 | 17 | 0.000 |
| **Inter-Gonial Width** | | | | | |
| Single Measures | .97 | 0.89 | 0.99 | 66 | 0.000 |
| Average Measures | .98 | 0.94 | 1.00 | 66 | 0.000 |
| **Anteroposterior Molar Length** | | | | | |
| Single Measures | .92 | 0.52 | 0.99 | 23 | 0.005 |
| Average Measures | .96 | 0.68 | 2.00 | 23 | 0.005 |
| **Root Length** | | | | | |
| Single Measures | .99 | 0.91 | 1 | 149 | 0.000 |
| Average Measures | .99 | 0.95 | 1 | 149 | 0.000 |
| **Molar Surface Area** | | | | | |
| Single Measures | .93 | 0.59 | 0.99 | 29 | 0.003 |
| Average Measures | .97 | 0.74 | 1.00 | 39 | 0.003 |
| **Incisor Anteroposterior Length** | | | | | |
| Single Measures | 0.96 | 0.7 | 1 | 55 | 0.009 |
| Average Measures | 0.98 | 0.87 | 1 | 55 | 0.009 |

### Supplemental Figure 3

**Cranial linear measurements in Chow, F1_HFHS, and F2_HFHS groups.** Quantitative assessment in male and female offspring from three experimental groups: Chow, F1_HFHS, and F2_HFHS. Measurements include skull length, midfacial length, skull width, alveolar bone length, neurocranium width, and inter-molar width (all in mm). Male data are represented in a blue gradient; female data are represented in a pink gradient. Values are presented as mean ± SEM. Statistical analysis was performed using one-way ANOVA; *p*-values shown above connecting lines indicate significance of intergroup comparisons.

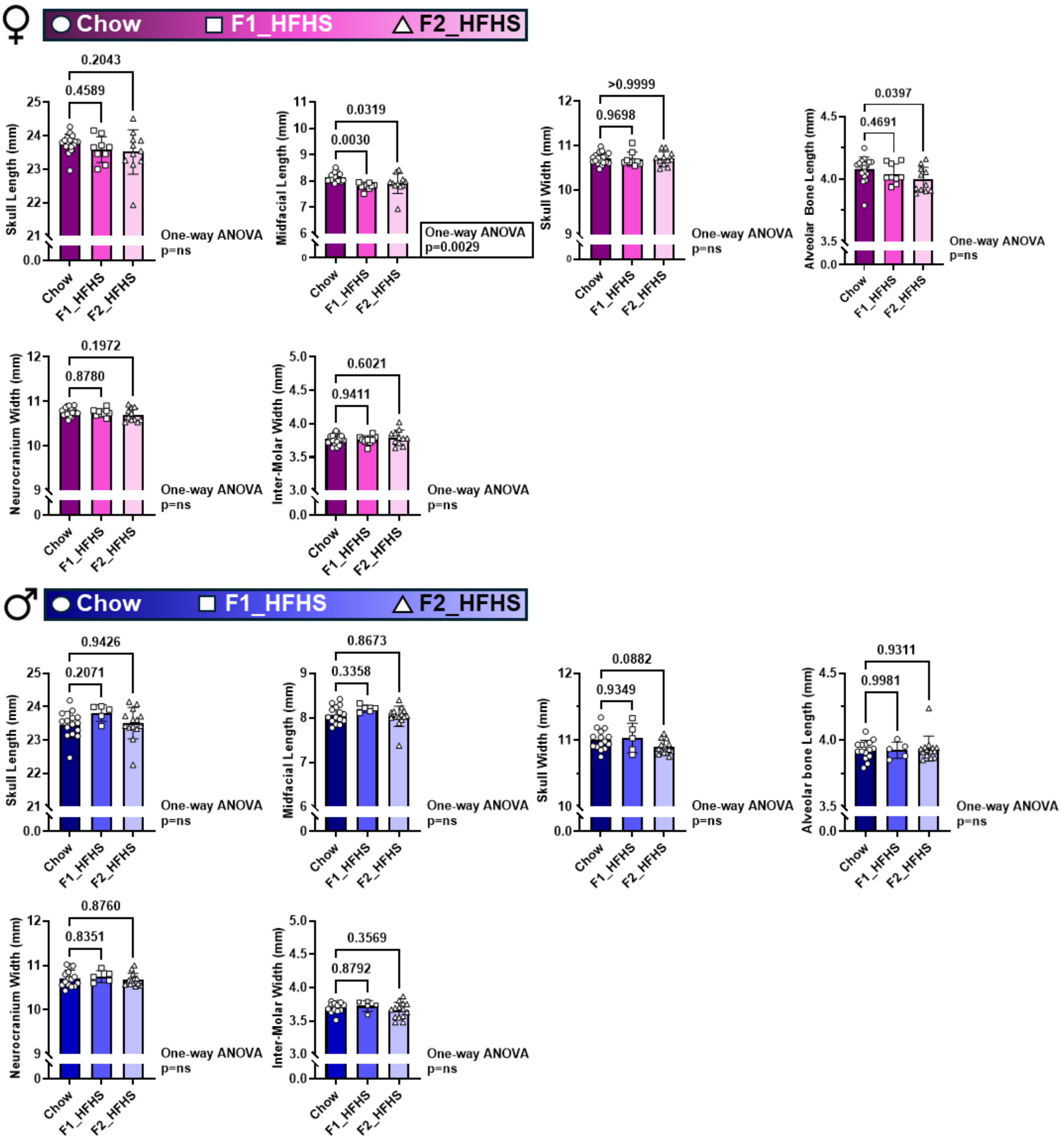

**Supplemental Figure 4**

**Mandible linear measurements in Chow, F1_HFHS, and F2_HFHS groups.** Quantitative assessment in male and female offspring from three experimental groups: Chow, F1_HFHS, and F2_HFHS. Measurements include condylar mandibular length, gonial mandibular length, mandibular height, inter-condylar width, alveolar bone length, inter-molar width, and inter-gonial width (all in mm). Male data are represented in a blue gradient; female data are represented in a pink gradient. Values are presented as mean ± SEM. Statistical analysis was performed using one-way ANOVA; *p*-values shown above connecting lines indicate significance of intergroup comparisons.

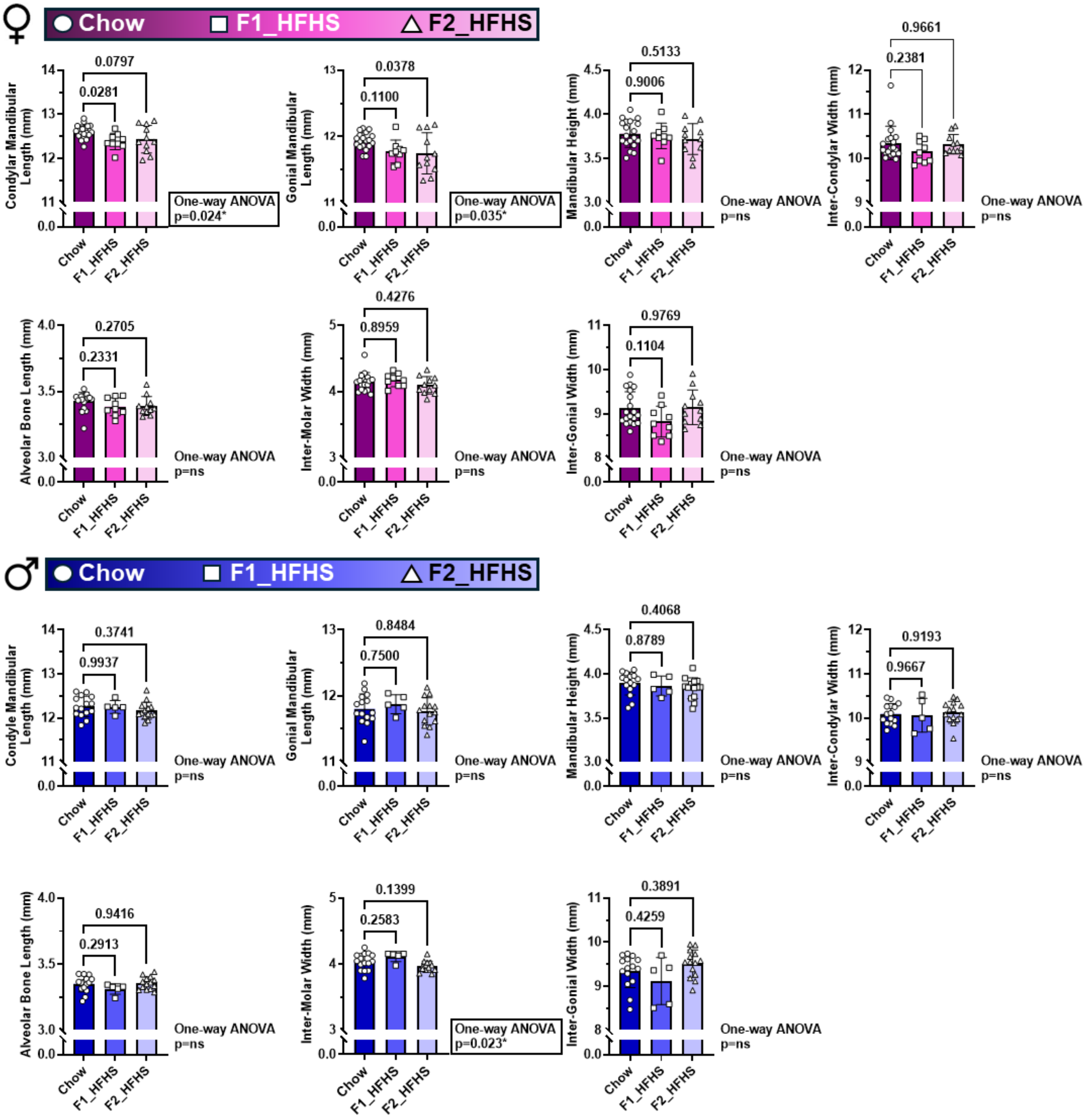

**Supplemental Figure 4**

**Dental measurements in Chow, F1_HFHS, and F2_HFHS groups.** Quantitative assessment in male and female offspring from three experimental groups: Chow, F1_HFHS, and F2_HFHS. Measurements include Anteroposterior molar length, Root length, crown surface area and incisor length  (all in mm). Male data are represented in a blue gradient; female data are represented in a pink gradient. Values are presented as mean ± SEM. Statistical analysis was performed using one-way ANOVA; *p*-values shown above connecting lines indicate significance of intergroup comparisons.

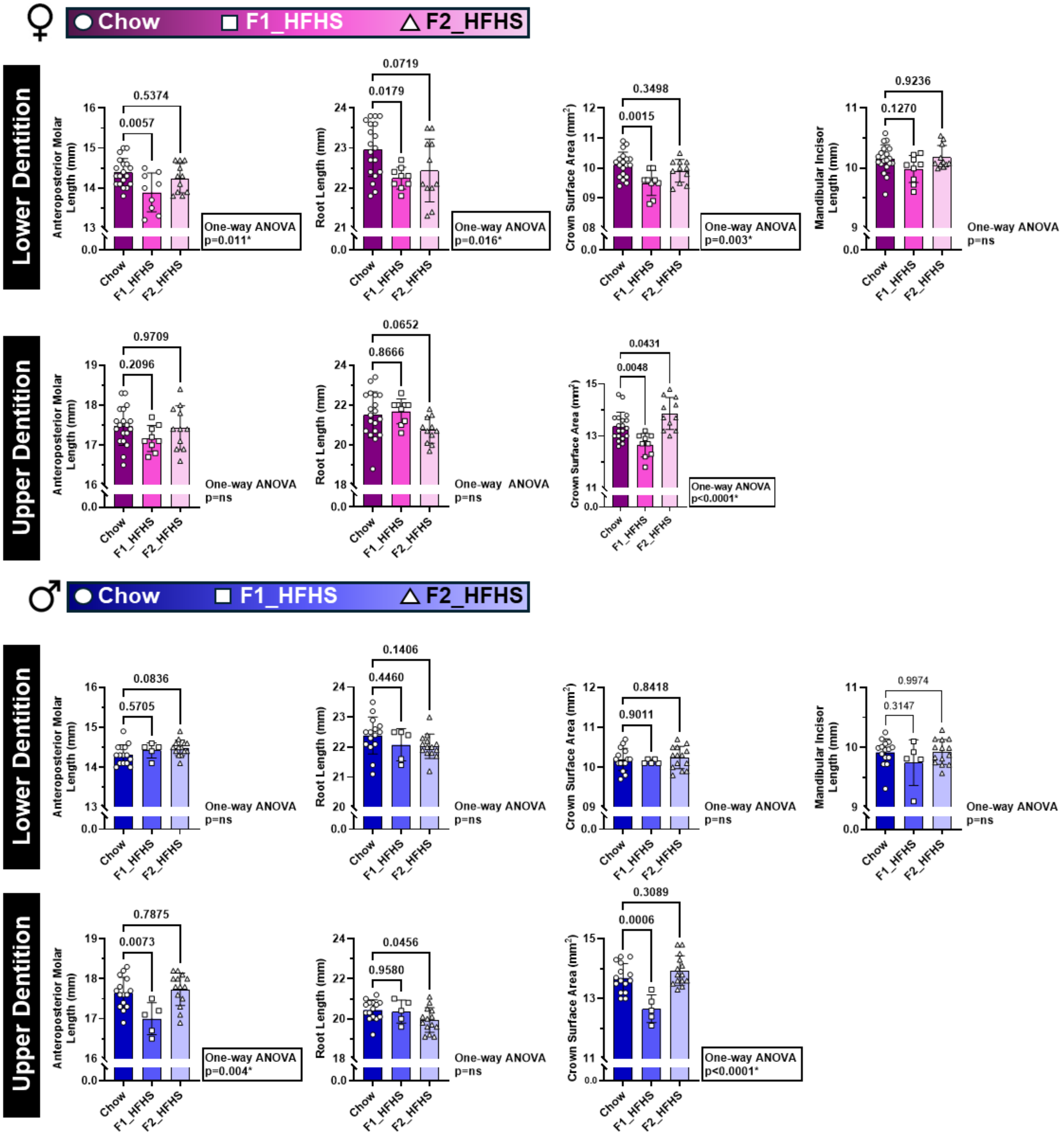

**Supplemental Figure 6**

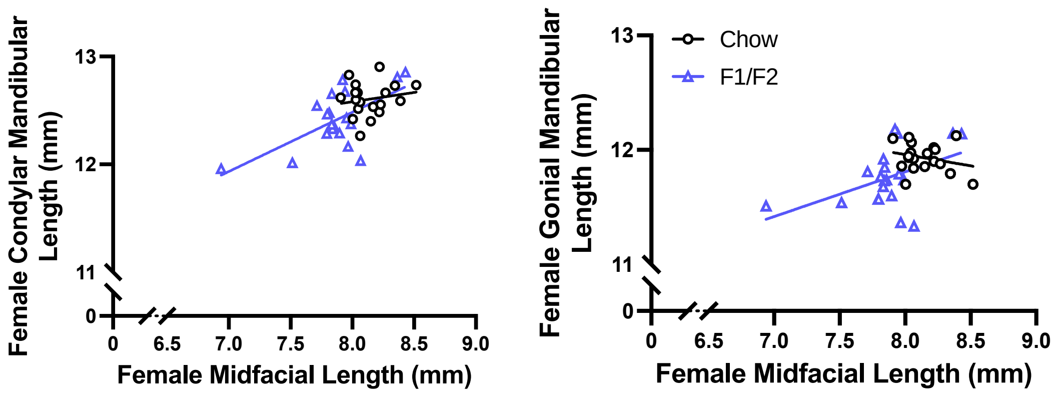


**Female Condyle and Gonial Correlation to Midfacial Length. Blue triangles represent F1 and F2. Black circles represent Chow. Significant correlation is observed.**

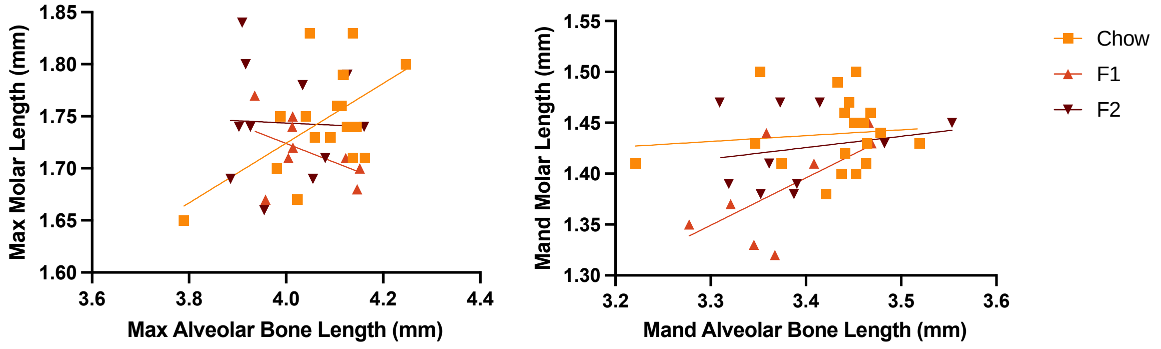


**Upper and lower Molar Length and corresponding alveolar bone length Correlation. No correlation is observed.**
